# Quantifying Somatic Mutation Burden: An Assay Validation Framework and Implementation in SomaticCODEC

**DOI:** 10.64898/2026.04.30.722079

**Authors:** Joshua N. Johnstone, James Phie, Cameron Fraser

## Abstract

Validation of somatic mutation burden assays is fundamentally constrained by the absence of a robust ground truth, limiting the interpretability of performance metrics. To address this, we propose a framework based primarily on relative validation, complemented by a suite of secondary metrics aligned to common failure modes. We implement this approach in SomaticCODEC, a ready-to-run assay for quantifying SNV burden in primary human samples, demonstrating strong linearity across mixtures of sperm and blood samples (R^2^ = 0.91) and high intra-batch precision (CV = 3.3%). This framework provides a practical approach for validating somatic mutation burden assays without requiring a ground truth.

## Introduction

Somatic mutations are analysed using a range of assays to address different scientific questions. These assays typically serve one of four analytical objectives. First, burden (also referred to as mutation load) quantifies the total number of somatic mutations at a given time, often for comparison across age or disease state. Second, composition (mutation spectrum or signature) describes the relative distribution of mutation types, providing insight into underlying mutational processes. Third, detection (variant identification or variant calling) focuses on identifying specific mutations, such as known resistance variants. Fourth, ranking (e.g. identification of top or enriched mutations) orders mutations by frequency, often to prioritise candidate driver events.

Measurement of somatic mutation burden is central to ageing and cancer research.Somatic mutations accumulate with age in most, if not all, human tissues (Ren, Dong and Vijg, 2022), and total mutation burden is associated with cancer risk through increased opportunities for clonal expansion (Martincorena and Campbell, 2015). In contrast to detection and ranking, which primarily capture high variant allele frequency (VAF) mutations, unbiased estimation of mutation burden requires sensitivity across the full VAF spectrum, including mutations present in only one or a few cells. Detection of low VAF mutations is challenging due to the background error rate of typical next-generation sequencing (NGS), which is several orders of magnitude higher than the true somatic mutation burden, particularly in healthy tissues (Kinde *et al*., 2011; Ren, Dong and Vijg, 2022).

Recent advances have substantially improved the feasibility of measuring somatic mutation burden. Central to these is duplex sequencing (Schmitt *et al*., 2012), which exploits the double-stranded nature of DNA to distinguish true mutations from most library preparation and sequencing errors that typically only occur on one strand. Further reductions in error rates have been achieved through improved library preparation methods, including fragmentation protocols that avoid error-prone end-repair and the use of specialised nucleotides to mitigate nick extension during A-tailing (Abascal *et al*., 2021). Early implementations of these approaches were limited by low throughput and restricted genomic coverage, reflecting the difficulty of ensuring both strands of a duplex molecule are sequenced. More recent methods have addressed this through strategies such as bottleneck dilution (Hoang *et al*., 2016) and physical concatenation of duplex strands (Bae *et al*., 2023), enabling broader and more efficient genome coverage.

The performance of any assay must be characterised to determine whether it is suitable for its intended use. Two validation paradigms are commonly applied. Absolute validation assesses accuracy against a reference or ground truth, whereas relative validation evaluates whether an assay can consistently differentiate relative differences between samples. Absolute validation requires either a reference standard of known quantity or a proxy ground truth. The validity of any proxy depends entirely on whether it accurately represents the true biological value.

A central challenge in validating somatic mutation burden assays is the absence of a reliable ground truth. With current technology, it is not possible to create a reference standard with a sufficiently low error rate: DNA synthesis and next-generation sequencing introduce errors at approximately 10^−5^ to 10^−2^ per base (Eckert and Kunkel, 1991; Carr *et al*., 2004; Kinde *et al*., 2011; Filges, Mouhanna and Ståhlberg, 2021), whereas somatic mutations occur at substantially lower frequencies, on the order of 10^−8^ to 10^−6^ per base depending on tissue (Ren, Dong and Vijg, 2022).

A commonly used approach is to use sperm as a proxy ground truth. This relies on the assumption that the mutation burden in sperm is close to zero, based on estimates from trio studies (Kong *et al*., 2012; Rahbari *et al*., 2016). However, trio studies estimate mutations transmitted to offspring, reflecting the highly selected subset of sperm that successfully contribute to fertilisation following an extreme selection bottleneck (∼1 in 10^8^–10^9^) (Cooper *et al*., 2009). It is therefore unlikely that these estimates are representative of the average mutation burden across sperm in a semen sample. In addition, comparison to a near-zero reference evaluates only an assay’s ability to avoid false positives and provides no information on sensitivity. Many studies combine these into a single “error rate”, obscuring the trade-off between sensitivity and specificity. Optimising against such a metric can bias assays toward high specificity with an unknown, and potentially severe, reduction in sensitivity.

Another commonly used approach is benchmarking against germline variant truth sets, such as those created by the Genome In A Bottle consortium (Zook *et al*., 2019). However, these are not suitable for assessing the performance of somatic mutation burden assays due to a fundamental mismatch in VAF. Germline variants typically occur at high VAF (approximately 50% or 100%), whereas somatic mutations relevant to burden occur at substantially lower biological frequencies (10^−8^ to 10^−6^) and are often present in only a small number of cells (Ren, Dong and Vijg, 2022). As a result, performance on germline truth sets does not reflect an assay’s ability to detect somatic mutations across the full VAF spectrum required for reliable burden estimation.

A third common approach is to benchmark performance against previous studies. While this can provide useful context, relying on it as a primary method of validation is problematic, as it constitutes a form of circular reasoning: agreement with prior estimates does not establish correctness in the absence of a ground truth.

To address the shortcomings of existing approaches to validating somatic mutation burden assays, we propose a general validation framework based primarily on relative validation (linearity and precision). This circumvents the challenges of establishing a reliable ground truth by removing the requirement for one. Under this framework, assays can continue to report absolute mutation burden, but their performance is assessed based on their ability to resolve relative differences between samples. For many use cases, this is sufficient; for example, a 20% reduction in SNV burden is likely to be biologically meaningful regardless of the absolute value. The utility of relative changes as a basis for interpretation is well established in other domains, including RNA-seq and quantitative PCR (Livak and Schmittgen, 2001; Wang, Gerstein and Snyder, 2009).

In addition, we define a suite of secondary validation metrics to complement relative validation by identifying failure modes not captured by linearity and precision. These include sentinel metrics of common failure modes, external concordance, and software validation.

Finally, we present SomaticCODEC, a ready-to-run assay for measuring single nucleotide variants (SNVs) in primary human samples from healthy donors, and evaluate its performance using this framework.

## Results and discussion

### Primary validation framework

The purpose of the primary validation metrics is to characterise an assay’s ability to perform relative quantitation of somatic mutation burden. This is captured by two metrics: linearity and precision.

Linearity assesses whether an assay produces a proportional response to changes in mutation burden within the tested range. It establishes that the measurement system preserves relative differences between samples, enabling meaningful comparison of magnitudes and interpolation between tested levels. However, linearity does not guarantee absolute accuracy: measurements may be incorrectly scaled or systematically biased.

Linearity is assessed by mixing samples with differing mutation burdens from the same donor. A low- and high-burden sample are combined in defined ratios, and the resulting measurements are evaluated for a proportional response. The choice of samples should reflect the range of mutation burdens relevant to the intended use case. For SomaticCODEC, we selected blood and sperm, as these are relevant to primary samples from healthy donors and are non-invasive to collect (Figure 1a). For other applications, alternative mixtures may be more appropriate, such as tumour and matched normal samples, or cultured cells with differing levels of mutagen exposure. We propose a minimum linearity threshold of R^2^ ≥ 0.8, defined on the lower bound of the 95% confidence interval (simple linear regression), with an ideal target of R^2^ ≥ 0.9.

**Figure 1.**
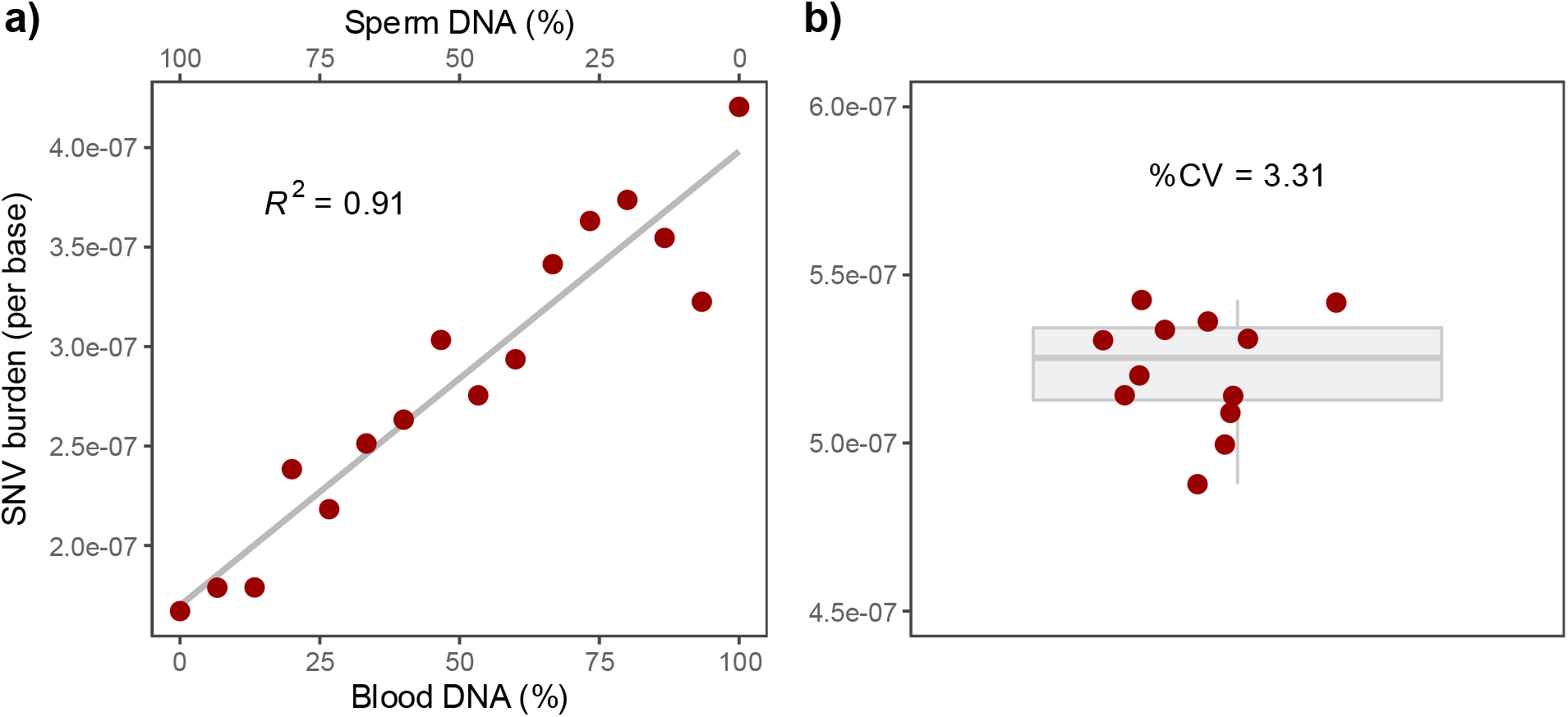
Primary validation of the SomaticCODEC assay. The performance of SomaticCODEC for relative quantification of SNV burden was characterised. **a)** Linearity was assessed by mixing DNA from blood and sperm at defined ratios; these tissues have relatively high and low SNV burdens, respectively. SomaticCODEC measured a linear increase in SNV burden with increasing proportions of blood DNA (N = 16, R^2^ = 0.91; one-sided lower 95% CI: 0.82, P < 1×10^-3^). **b)** Intra-batch precision was measured by processing 12 replicates of the same buffy coat sample in a single library preparation and sequencing batch. The coefficient of variation (CV) was 3.31% (One-sided upper 95% CI: 4.02%). Each red point represents one individual sample.

Precision assesses the consistency of the measurement system across repeated measurements. It can be evaluated as intra-batch precision (within a single batch) or inter-batch precision (across batches). Reproducibility extends this further to consistency across laboratories or research groups.

We recommend quantifying precision using either the coefficient of variation (CV) or normalized interquartile range (norm-IQR). For intra-batch precision (Figure 1b), we propose a minimum threshold of CV ≤ 20%, defined on the upper bound of the 95% confidence interval, with an ideal target of ≤ 10%. We recommend a minimum of 12 samples to estimate the confidence interval, as bootstrap-based intervals can be unreliable at smaller sample sizes (Chernick, 2008).

### Secondary validation framework

While relative validation should form the primary assessment of a somatic mutation burden assay, it does not capture all potential failure modes. We therefore introduce a set of secondary validation metrics designed to identify gaps that are not revealed by linearity and precision alone. These metrics are not intended to provide definitive measures of performance in isolation; rather, they serve as indicators that can highlight potential issues and prompt further investigation. We group these into three categories: sentinels of common failure modes, external concordance, and software testing.

### Sentinels of common failure modes

Sentinels of common failure modes comprise a diverse set of metrics that assess assay performance across the full workflow, from library preparation through to variant calling. For SomaticCODEC, we implemented a comprehensive set of 91 metrics (Supplementary Table 1), described as component-level and system-level metrics, which are automatically reported for every sample against predetermined thresholds. In the main text, we discuss a subset of these metrics, but we recommend adoption of the full set.

Germline leakage is a pervasive challenge for somatic mutation assays, as germline variants occur at substantially higher frequencies than somatic variants (Trost, Loureiro and Scherer, 2021). Even low rates of misclassification can result in a large proportion of false positive calls in the somatic variant callset. These variants may originate from the same donor (incomplete masking of germline variants) or from another donor (DNA contamination). For SomaticCODEC, we assess germline leakage by comparing called variants to a subset of the gnomAD v4 database (AF ≥ 1 × 10^−3^; Figure 2a). Overlap with gnomAD does not by itself indicate germline leakage, as some overlap is expected due to shared mutational processes and the large size of the dataset. This metric is most informative when interpreted relative to typical values, with elevated overlap identifying samples that may be contaminated.

**Figure 2.**
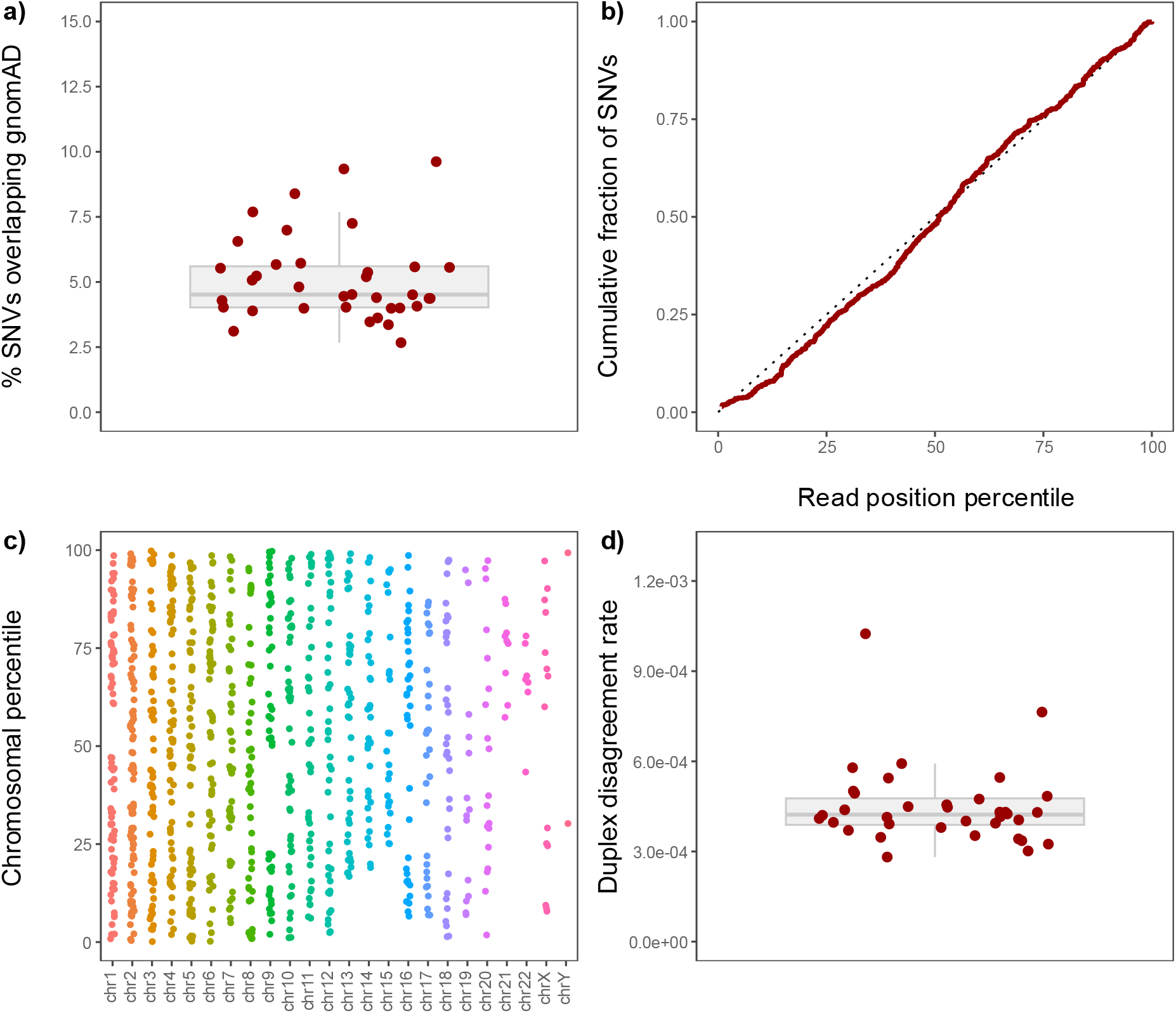
Detection of failure modes in SomaticCODEC. Four of the 91 metrics used to identify common failure modes. **a)** Overlap of called variants with the gnomAD v4 dataset (AF > 1×10^-3^). Across 36 buffy coat samples, a median of 4.52% (95% CI: 4.29–5.37%) of somatic SNVs overlapped with gnomAD. **b)** Distribution of SNVs along the read length. Across nine buffy coat samples, the median of the maximum absolute deviation from a uniform distribution was 4.7% (95% CI: 3.9–4.9%), indicating near-uniformity. The dotted line represents a uniform distribution, and the red line shows a representative sample. **c)** Distribution of SNV calls across the genome. Across 36 buffy coat samples, the median genomic distribution index was 11.91% (95% CI: 10.77–12.78%), indicating broad genomic coverage. See Supplementary Methods for definition. The genomic distribution of called SNVs is shown for a representative sample. **d)** Duplex disagreement rate for 36 buffy coat samples. The median was 4.23 × 10^-4^ (95% CI: 4.01 × 10^-4^–4.47 × 10^-4^). Under the assumption of symmetric duplex errors across strands and positions, this corresponds to an estimated error rate of 1.49 × 10^-8^.

Non-uniform distributions of called mutations by read position, such as clustering at read ends, may indicate technical artefacts rather than genuine biological mutations. For SomaticCODEC, SNV calls were distributed approximately uniformly across read positions (Figure 2b).

Somatic mutation burden assays assume representative sampling of the genome. Under-representation of specific genomic regions may indicate regional bias in the assay. Such biases can arise from enzymatic fragmentation during library preparation or imbalanced genome masking. For SomaticCODEC, SNVs were distributed across all chromosomes without evidence of meaningful regional bias (Figure 2c).

Duplex-based somatic mutation burden assays represent the current gold standard. The duplex disagreement rate quantifies the rate of disagreement between Watson and Crick strands during consensus calling. This is a powerful indicator of assay performance, as it captures a wide range of failure modes arising across library preparation and sequencing. Under the assumption of symmetric duplex errors across strands and positions, which may not hold in practice, the assay error rate can be estimated from the duplex disagreement rate. Under this assumption, the median duplex disagreement rate reported for SomaticCODEC of 4.23 × 10^-4^ (Figure 2d) corresponds to an estimated error rate of 1.49 × 10^-8^. Despite uncertainty in the absolute interpretation of this metric, substantial deviations from typical values can indicate regression at the level of individual samples or batches.

### External concordance

External concordance is useful for identifying gross discrepancies with prior work, but should not be relied upon as a primary means of validation or treated as a surrogate ground truth. Somatic SNV burden and composition measured by SomaticCODEC were compared with prior high-accuracy duplex sequencing assays in the same tissues.

SomaticCODEC estimates an association between buffy coat SNV burden and age with a slope of 4.2 × 10^-9^ (95% CI: 2.7 × 10^-9^–5.6 × 10^-9^; P < 0.001; Figure 3a), which is similar to slopes calculated from buffy coat data from Bae *et al*. (2023) (3.4 × 10^-9^) and granulocyte data from Abascal *et al*. (2021) (3.7 × 10^-9^).

**Figure 3.**
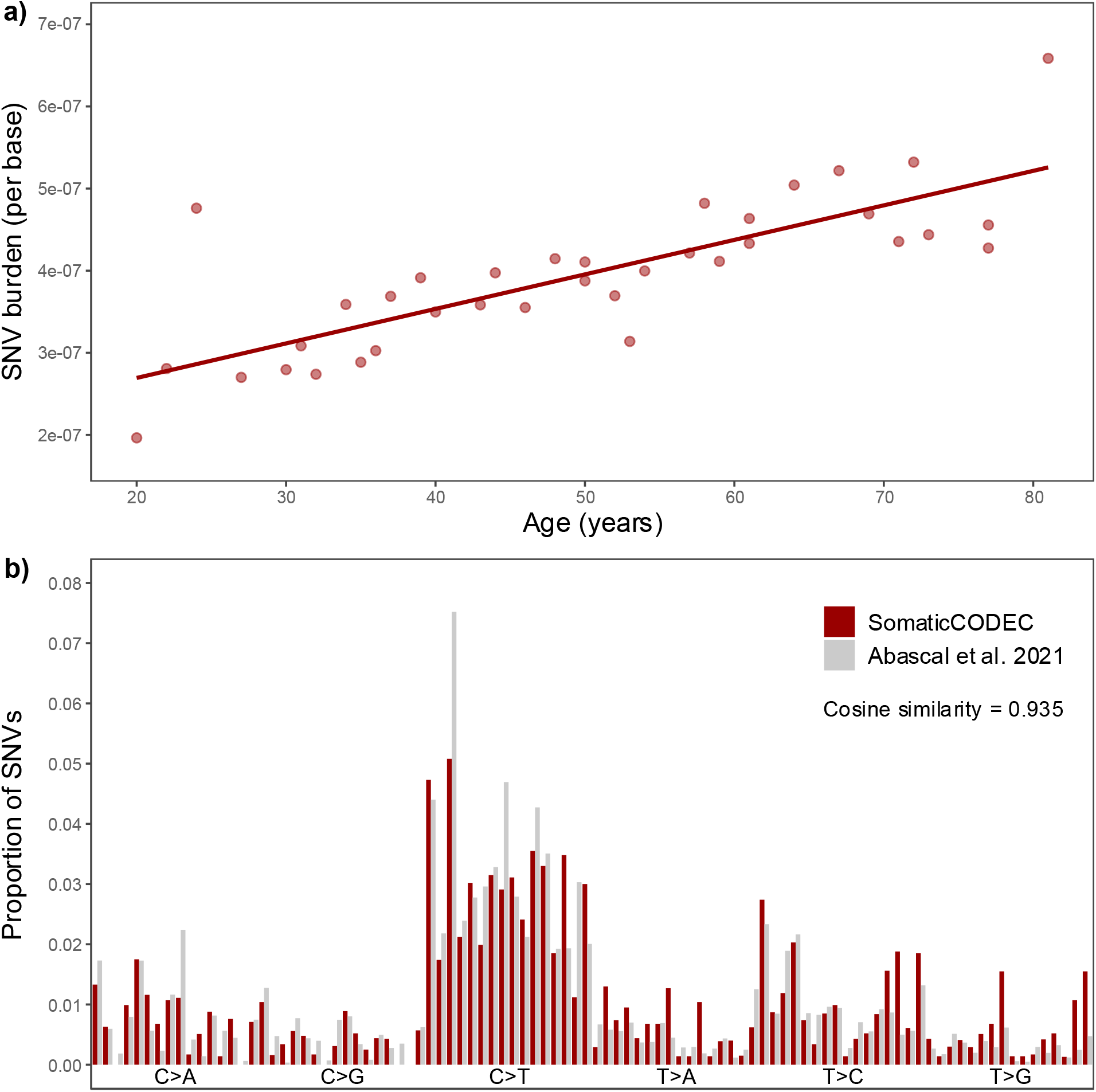
Concordance of SomaticCODEC with external data. The SNV burdens and compositions measured by SomaticCODEC. **a)** Association between buffy coat SNV burden and age measured by the SomaticCODEC assay (slope = 4.2 × 10^-9^, 95% CI = [2.7 × 10^-9^, 5.6 × 10^-9^], N = 36 *P* < 1×10^-3^). **b)** Trinucleotide signatures measured by SomaticCODEC (buffy coat) and Abascal *et al*. (2021) (granulocytes). Trinucleotide signatures from 12 buffy coat samples closely match those of Abascal *et al*. (2021), with a median cosine similarity of 0.934 (95% CI = [0.926, 0.948]). The red bars show a representative SomaticCODEC trinucleotide signature, while the grey bars show the signature reported by Abascal *et al*. (2021).

Trinucleotide context was compared between buffy coat samples from SomaticCODEC and granulocyte samples from Abascal *et al*. (2021). The median cosine similarity was 0.934 (95% CI: 0.926–0.948; Figure 3b), indicating highly similar trinucleotide profiles between studies. Deviations in trinucleotide context can highlight failure modes; for example, an elevated proportion of C→A variants in an early version of the SomaticCODEC library preparation revealed oxidative damage (Supplementary Figure 1).

Sperm SNV burden was compared to Abascal *et al*. (2021) and Shoag *et al*. (2025). SomaticCODEC measured higher sperm SNV burdens than Abascal *et al*. (2021) (P < 0.001, Tukey’s HSD) and Shoag *et al*. (2025) (P < 0.001, Tukey’s HSD) (Supplementary Figure 2). This discrepancy may reflect a higher true sperm SNV burden than previously reported, for example if prior studies applied filtering strategies that prioritise minimising error rates (specificity) at the expense of sensitivity, thereby suppressing true variants. Alternatively, it may arise from false positive mutation calls in SomaticCODEC that are not detected by our sentinel metrics.

### Software validation

Most somatic mutation burden assays rely on complex software pipelines to convert raw sequencing data into scientific outputs. Software errors can lead to incorrect mutation calls that may not be detected by other validation approaches. For example, incorrectly applied genome masking may not be apparent to operators but can meaningfully affect variant calling.

Ad hoc testing can detect some failure modes but is high effort and limited to the time of testing. Many errors arise when changes are made to the codebase or from interactions between components. Automated testing requires additional upfront effort but enables the software to be retested after each change at minimal cost, providing ongoing protection against regression.

For SomaticCODEC, we implemented 318 software tests using the pytest framework (Krekel *et al*., 2004). These tests use real sequencing data, reduced to a few thousand reads and a single chromosome to enable rapid execution. We implemented at least one unit test for each Snakemake (Mölder *et al*., 2021) rule, with multiple tests for higher-risk rules. These tests verify that each rule produces expected outputs from known inputs, substantially reducing the risk of undetected errors, particularly during code modification.

### Change control framework

Any change to an assay has the potential to alter performance. While changes are typically intended to improve performance, they may also introduce unintended effects. Once an assay has been validated to achieve a high level of performance, it is recommended practice to place controls on subsequent changes until their impact can be demonstrated to be at least performance-neutral or beneficial.

For SomaticCODEC, key components were subject to change control. Library preparation methods were defined in version-controlled SOPs, the bioinformatic pipeline was version controlled using Git, and the computational environment was containerised using Docker with pinned tool versions. These were only changed with evidence of net improvement in performance.

### SomaticCODEC assay

In addition to the validation framework, we developed an assay for measuring somatic SNV burden in healthy human tissue (SomaticCODEC). The following sections describe the design decisions underlying the development of the SomaticCODEC assay.

### Library preparation

As somatic variant calling in healthy tissues requires low error rates for accurate detection of ultra-low VAF mutations, we selected a library preparation workflow designed to minimise technical artefacts. Library preparation was performed using CODEC, a duplex sequencing method that enables identification of true duplex-supported variants while suppressing single-strand library preparation and sequencing errors (Bae *et al*., 2023). We further incorporated ddNTP-blocked A-tailing and blunt-end enzymatic fragmentation to reduce nick- and end-repair artefacts (Abascal *et al*., 2021).

### Unbiased estimation of SNV burden

Accurate estimation of SNV burden requires an unbiased relationship between the number of observed variants (numerator) and the number of genomic positions assessed (denominator). To achieve this, all filtering and masking of reads and bases were performed before SNV calling, ensuring they were applied independently of SNV status. Filtering reads after variant calling inherently leads to underestimation of SNV burden if the same filtering is not applied to all analysed bases.

### Genome masking

To reduce germline leakage from the individual’s own germline, we use a “germline risk” mask. While other somatic mutation assays identify germline mutations with callers such as GATK’s HaplotypeCaller (Van der Auwera and O’Connor, 2020), these callers require high confidence to call germline mutations. This allows low-confidence mutations, particularly heterozygous mutations with support for two different alleles, to leak into the somatic callset. SomaticCODEC addresses this issue by instead using a pileup-based approach to identify “germline risk” sites. For each position, the VAF for each allele is calculated across the reads. Any position with total alternate VAF exceeding a specific threshold (by default 10%) is added to the germline risk mask, greatly increasing the chance of masking germline SNVs. Positions with insufficient depth for germline risk calling are added to a separate low depth mask. A limitation of this approach is that high VAF somatic SNVs may be masked; however, given that most somatic SNVs will have ultra-low VAF, this is a reasonable trade-off.

To complement our germline risk masking, and to protect against germline leakage from inter-sample DNA contamination, we mask a subset of common germline variants from the gnomAD v4 database (Chen *et al*., 2024). Masking the full gnomAD dataset would substantially reduce sensitivity, given that it contains more than 900 million variants (Van Buren *et al*., 2025), and germline and somatic SNVs share a common mechanistic basis. We instead mask only variants with an allele frequency ≥ 0.1, reducing germline leakage from the most common variants while preserving sensitivity.

Misalignment of reads can produce artefactual SNV calls (Li, 2014; Bowler *et al*., 2019). To mitigate these artefacts, we masked various difficult-to-map genomic regions. We used RepeatMasker (Smit, Hubley and Green, 2015) to identify interspersed repeats and low complexity sequences in the reference genome. We also used the Genome In A Bottle consortium’s difficult regions mask (Olson *et al*., 2022), which includes segmental duplications, large homopolymers, and other difficult regions identified through variant call benchmarking (Wagner *et al*., 2022). In combination, these masks cover >50% of the genome; it may be possible to reduce the size of the mask and the resulting effect on sensitivity by further refining the regions most prone to misalignment.

### Matched sample design

A matched sample design uses a second sample from the same donor to identify germline variants, enabling the differentiation of high VAF somatic variants from true germline variants, especially if a different tissue is used. A further advantage is that matched samples can be sequenced using lower cost (non-duplex) approaches, as high-depth germline calling does not require duplex error correction. Our matched sample is fragmented using the same method as the CODEC sample to maximise overlap between samples.

### Variant calling

Somatic SNV burden was measured using a calling framework that permits variants to be accepted even when observed only once, provided they are supported by high-confidence duplex consensus data. This design reflects the intended use of the assay, which requires quantification of mutation burden in healthy tissues where SNVs are ultra-low VAF and may be private to only one or a few cells. Increased confidence in a variant call is derived primarily from per-molecule base-calling accuracy rather than repeated observation through sequencing depth.

This differs from most conventional somatic variant callers, such as Mutect2 (Van der Auwera and O’Connor, 2020), which were developed largely for cancer-focused applications involving detection and frequency estimation of specific mutations. In those settings, confidence that a true variant exists, and estimation of its VAF, are strengthened by multiple independent observations of the alternate allele, and single observations are rejected due to low confidence. Applying the same depth-based framework to burden analysis in healthy tissue would preferentially exclude ultra-low VAF mutations and bias mutation burden estimates downward.

We therefore used a pileup-based workflow with bcftools mpileup (Danecek *et al*., 2021) to enable transparent counting of duplex-supported somatic SNV observations at each analysable position. Somatic SNV burden was calculated as the number of SNVs detected across unmasked, analysable positions divided by the total number of unmasked, analysable positions. Base level filtering focused on properties relevant to individual call reliability, including base quality, masking of difficult regions, and masking positions with low germline depth. The base quality threshold was set to Q70, as including bases below this threshold is likely to result in an unacceptable false positive SNV rate when calling variants using this method.

## Methods

DNA was extracted from thawed buffy coat, blood and lysed sperm using the Qiagen DNA Mini Kit, following the manufacturer’s protocol with minor modifications. Library preparation and sequencing were performed as described in Phie *et al*. (2026), with minor modifications. Data processing and somatic SNV calling were performed using SomaticCODEC v5.0.0 (Johnstone, Phie and Fraser, 2026). Statistical analyses were conducted in R v4.3 (R Core Team, 2021). Detailed methods are provided in the Supplementary Information.

## Author contributions

Joshua Johnstone: Methodology, Software, Validation, Formal Analysis, Investigation, Writing – Original Draft, Writing – Review & Editing.

James Phie: Methodology, Software, Formal Analysis, Investigation, Writing – Original Draft, Writing – Review & Editing.

Cameron Fraser: Conceptualization, Methodology, Software, Validation, Formal Analysis, Writing – Original Draft, Writing – Review & Editing, Supervision, Project Administration.

## Data availability

Raw sequencing data are subject to ethical and regulatory restrictions and cannot be made publicly available at this time. Access may be possible through controlled-access repositories (e.g. dbGaP) subject to appropriate approvals. The availability of raw data is dependent on institutional and regulatory approvals. Processed data supporting the findings of this study are available from the authors upon reasonable request.

## Code availability

A ready-to-run bioinformatics pipeline is available at https://github.com/systematicmedicine/SomaticCODEC. All analyses in this study were performed using version v5.0.0. A more recent version (v6.0.2) is also available, which includes additional quality-of-life improvements.

## Acknowledgements

We thank Scott Needham and Leading Technology Group for funding this work and supporting its dissemination to the broader research community. We also acknowledge Scott Needham’s contribution to project conceptualisation.

We thank Fulcrum Genomics Pty Ltd for their advice on duplex variant calling assays, and Jin Bae for his advice on CODEC library preparation methodology.

We are grateful to all donors who provided samples used in this research.

## References

Abascal, F. et al. (2021) “Somatic mutation landscapes at single-molecule resolution,” Nature, 593(7859), pp. 405–410. Available at: 10.1038/s41586-021-03477-4.

Bae, J.H. et al. (2023) “Single duplex DNA sequencing with CODEC detects mutations with high sensitivity,” Nature Genetics, 55(5), pp. 871–879. Available at: 10.1038/s41588-023-01376-0.

Bowler, T.G. et al. (2019) “Misidentification of MLL3 and other mutations in cancer due to highly homologous genomic regions,” Leukemia & Lymphoma, 60(13), pp. 3132–3137. Available at: 10.1080/10428194.2019.1630620.

Carr, P.A. et al. (2004) “Protein-mediated error correction for de novo DNA synthesis,” Nucleic Acids Research, 32(20), pp. e162–e162. Available at: 10.1093/nar/gnh160.

Chen, S. et al. (2024) “A genomic mutational constraint map using variation in 76,156 human genomes,” Nature, 625(7993), pp. 92–100. Available at: 10.1038/s41586-023-06045-0.

Chernick, M.R. (2008) “Bootstrap Methods: A Guide for Practitioners and Researchers,” Biometrics, 64(3), pp. 998–999. Available at: 10.1111/j.1541-0420.2008.01082_17.x.

Cooper, T.G. et al. (2009) “World Health Organization reference values for human semen characteristics,” Human Reproduction Update, 16(3), pp. 231–245. Available at: 10.1093/humupd/dmp048.

Danecek, P. et al. (2021) “Twelve years of SAMtools and BCFtools,” GigaScience, 10(2), p. giab008. Available at: 10.1093/gigascience/giab008.

Eckert, K.A. and Kunkel, T.A. (1991) “DNA Polymerase Fidelity and the Polymerase Chain Reaction,” PCR Methods and Applications [Preprint]. Available at: 10.1101/gr.1.1.17.

Filges, S., Mouhanna, P. and Ståhlberg, A. (2021) “Digital Quantification of Chemical Oligonucleotide Synthesis Errors,” Clinical Chemistry, 67(10), pp. 1384–1394. Available at: 10.1093/clinchem/hvab136.

Hoang, M.L. et al. (2016) “Genome-wide quantification of rare somatic mutations in normal human tissues using massively parallel sequencing,” Proceedings of the National Academy of Sciences, 113(35), pp. 9846–9851. Available at: 10.1073/pnas.1607794113.

Johnstone, J.N., Phie, J. and Fraser, C. (2026) SomaticCODEC. Systematic Medicine. Available at: https://github.com/systematicmedicine/SomaticCODEC.

Kinde, I. et al. (2011) “Detection and quantification of rare mutations with massively parallel sequencing,” Proceedings of the National Academy of Sciences, 108(23), pp. 9530–9535. Available at: 10.1073/pnas.1105422108.

Kong, A. et al. (2012) “Rate of de novo mutations and the importance of father’s age to disease risk,” Nature, 488(7412), pp. 471–475. Available at: 10.1038/nature11396.

Krekel, H. et al. (2004) pytest 8.4. Available at: https://github.com/pytest-dev/pytest.

Li, H. (2014) “Toward better understanding of artifacts in variant calling from high-coverage samples,” Bioinformatics, 30(20), pp. 2843–2851. Available at: 10.1093/bioinformatics/btu356.

Livak, K.J. and Schmittgen, T.D. (2001) “Analysis of Relative Gene Expression Data Using Real-Time Quantitative PCR and the 2™ΔΔC T Method,” Methods, 25(4), pp. 402–408. Available at: 10.1006/meth.2001.1262.

Martincorena, I. and Campbell, P.J. (2015) “Somatic mutation in cancer and normal cells,” Science, 349(6255), pp. 1483–1489. Available at: 10.1126/science.aab4082.

Mölder, F. et al. (2021) “Sustainable data analysis with Snakemake,” F1000Research, 10, p. 33. Available at: 10.12688/f1000research.29032.2.

Olson, N.D. et al. (2022) “PrecisionFDA Truth Challenge V2: Calling variants from short and long reads in difficult-to-map regions,” Cell Genomics, 2(5), p. 100129. Available at: 10.1016/j.xgen.2022.100129.

Phie, J. et al. (2026) “CODEC Library Preparation From Genomic DNA,” Under review [Preprint].

Rahbari, R. et al. (2016) “Timing, rates and spectra of human germline mutation,” Nature Genetics, 48(2), pp. 126–133. Available at: 10.1038/ng.3469.

R Core Team (2021) R: A language and environment for statistical computing. Vienna, Austria: R Foundation for Statistical Computing. Available at: https://www.R-project.org/.

Ren, P., Dong, X. and Vijg, J. (2022) “Age-related somatic mutation burden in human tissues,” Frontiers in Aging, 3, p. 1018119. Available at: 10.3389/fragi.2022.1018119.

Schmitt, M.W. et al. (2012) “Detection of ultra-rare mutations by next-generation sequencing,” Proceedings of the National Academy of Sciences, 109(36), pp. 14508–14513. Available at: 10.1073/pnas.1208715109.

Shoag, J.E. et al. (2025) “Direct measurement of the male germline mutation rate in individuals using sequential sperm samples,” Nature Communications, 16(1), p. 2546. Available at: 10.1038/s41467-025-57507-0.

Smit, A., Hubley, R. and Green, P. (2015) RepeatMasker Open-4.0. Available at: http://www.repeatmasker.org.

Trost, B., Loureiro, L.O. and Scherer, S.W. (2021) “Discovery of genomic variation across a generation,” Human Molecular Genetics, 30(20), pp. R174–R186. Available at: 10.1093/hmg/ddab209.

Van Buren, S.L. et al. (2025) “A comparative review of short genetic variant databases across humans and animal species,” Briefings in Bioinformatics, 26(4), p. bbaf356. Available at: 10.1093/bib/bbaf356.

Van der Auwera, G.A. and O’Connor, B.D. (2020) Genomics in the Cloud: Using Docker, GATK, and WDL in Terra. 1st ed. O’Reilly Media.

Wagner, J. et al. (2022) “Benchmarking challenging small variants with linked and long reads,” Cell Genomics, 2(5), p. 100128. Available at: 10.1016/j.xgen.2022.100128.

Wang, Z., Gerstein, M. and Snyder, M. (2009) “RNA-Seq: a revolutionary tool for transcriptomics,” Nature Reviews Genetics, 10(1), pp. 57–63. Available at: 10.1038/nrg2484.

Zook, J.M. et al. (2019) “An open resource for accurately benchmarking small variant and reference calls,” Nature Biotechnology, 37(5), pp. 561–566. Available at: 10.1038/s41587-019-0074-6.

