## Supplementary Information for "Quantifying Somatic Mutation Burden: An Assay Validation Framework and Implementation in SomaticCODEC"

#### Supplementary results

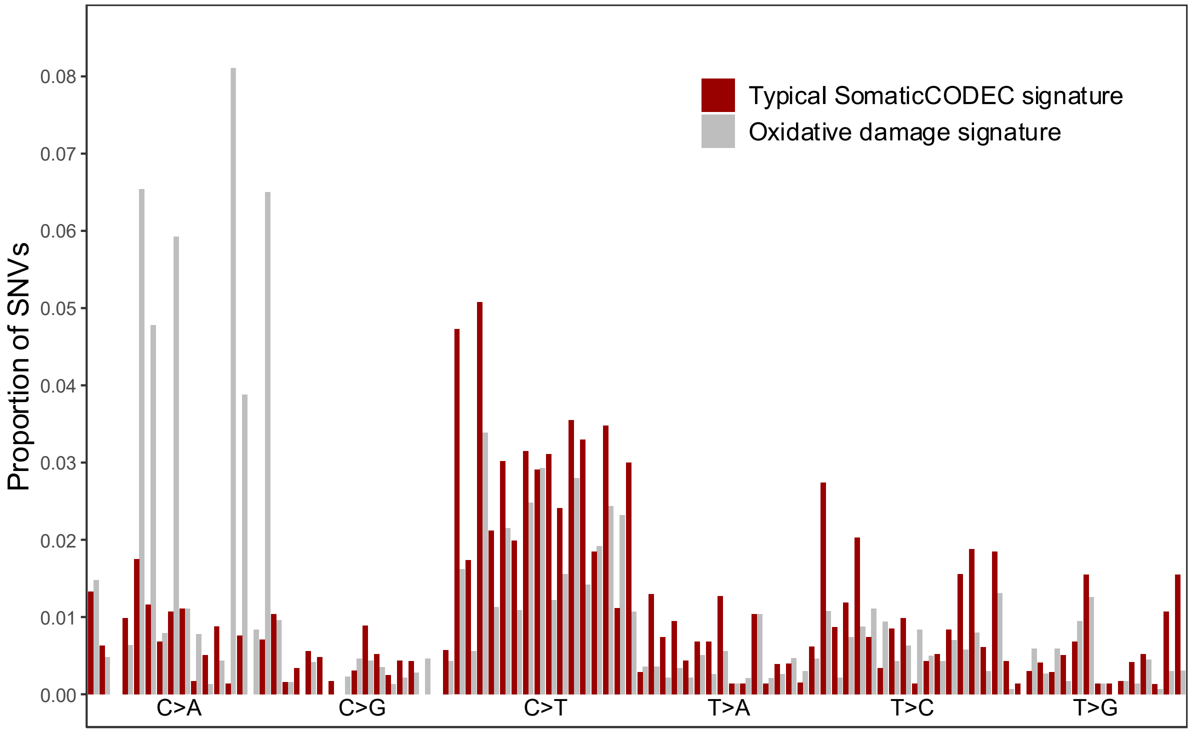


**Supplementary Figure 1 – Trinucleotide signature with evidence of oxidative damage.** Trinucleotide signatures can be compared to detect potential library preparation and sequencing artefacts. The red bars show a typical SomaticCODEC buffy coat trinucleotide signature, while the grey bars show a signature from a sample with an elevated proportion of C>A variants; indicative of oxidative damage.

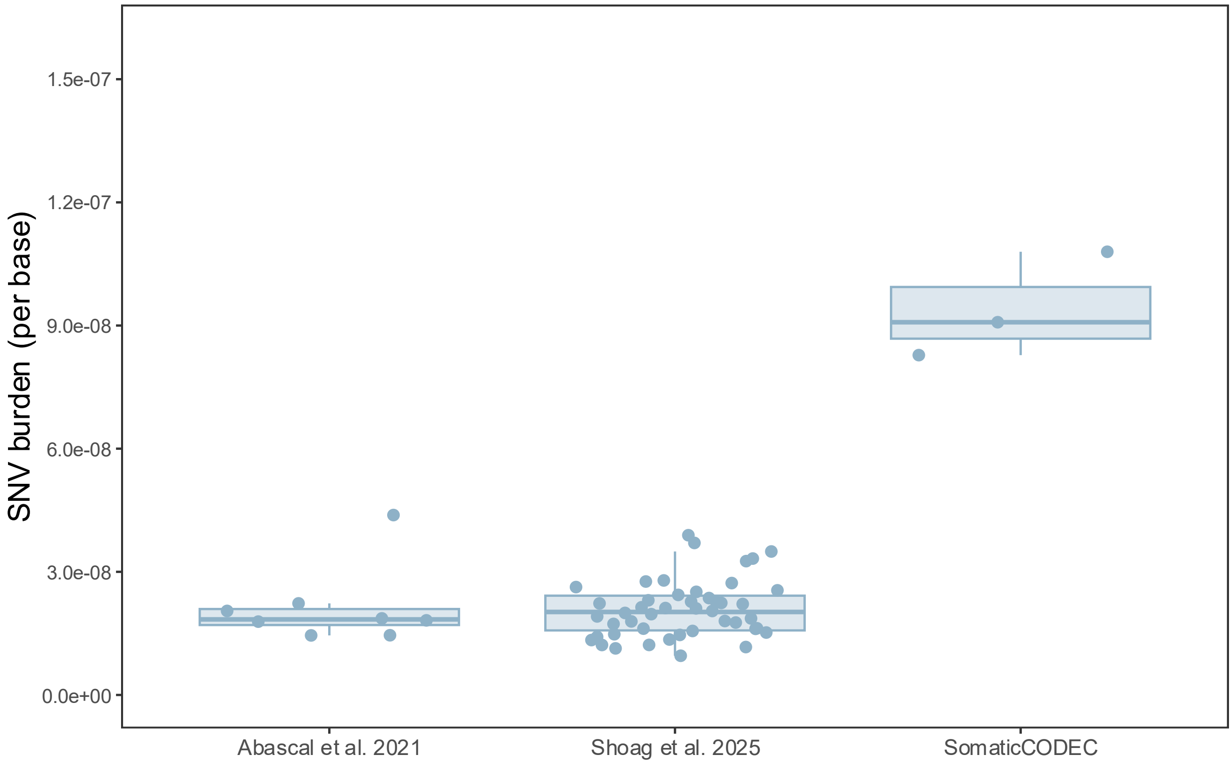


**Supplementary Figure 2 – External concordance for sperm SNV burden.**

Sperm somatic SNV burdens as reported by external studies and SomaticCODEC. The mean burden measured by SomaticCODEC (9.39 × 10-8, 95% CI = [$6.19 \times 10^{-8}$, $1.26 \times 10^{-7}$]) is higher than burdens reported by Abascal *et al*. (2021) (Mean = $2.13 \times 10^{-8}$, 95% CI = [1.34 × 10-8, $2.92 \times 10^{-8}$], Tukey's HSD: P < 0.001) and Shoag *et al*. (2025) (Mean = $2.08 \times 10^{-8}$, 95% CI = [$1.88 \times 10^{-8}$, $2.29 \times 10^{-8}$], Tukey's HSD: P < 0.001).

### Supplementary methods

#### Ethics Approval

This study was approved by the Human Research Ethics Committee of St Vincent's Hospital Melbourne (HREC 130/22). Written informed consent was obtained from all participants. Remnant samples were provided by Australian Red Cross Lifeblood under appropriate ethical and institutional approvals.

#### Sample Collection

Remnant buffy coats from a random subset of blood donors in the Melbourne, Australia area were supplied by the Australian Red Cross Lifeblood service. Whole blood samples (36 mL) were collected in lithium heparin tubes by phlebotomists at Emeritus Research. Semen samples were collected by Monash IVF Pty Ltd.

#### DNA extraction

##### Buffy coat and whole blood DNA extraction

DNA was extracted from frozen whole blood and buffy coat using a QIAamp DNA Mini Kit (Qiagen Cat. #51304). The kit manual was followed with minor modifications: all centrifugation steps were performed at 20000 x g, and elution was performed with 100 µL of LTE buffer (10mM Tris-HCl pH 8, 0.1mM EDTA) for whole blood samples, or 200 µL of LTE buffer for buffy coat samples. The eluted DNA was stored at -20°C in 1.5 mL LoBind tubes (Eppendorf Cat. #30108051) until use.

##### Sperm lysis and DNA extraction

Cryopreserved semen samples were thawed by swirling the straws in a 37°C water bath until just thawed. A 15 mL centrifuge tube was prepared for density gradient separation by adding 1 mL ORIGIO Gradient 80 medium (Cooper Surgical Cat. #84022060), followed by 1 mL ORIGIO Gradient 40 medium (Cooper Surgical Cat. #84022060). 1 mL of semen was added above the gradient 40 layer, and the tube was centrifuged at 400 x g for 15 minutes. The supernatant was removed from the pellet and discarded. In a new 15 mL centrifuge tube, 5 mL of ORIGIO Sperm Wash (Cooper Surgical Cat. #84055060) was added, then the pellet was transferred to this tube and pipette mixed with the Sperm Wash. The tube was centrifuged at 300 x g for 5 minutes. The supernatant was removed and discarded, then 5 mL Sperm Wash was added to the tube and pipette mixed. The tube was again centrifuged at 300 x g for 5 minutes, then the supernatant was removed and discarded, leaving ~50 µL of solution at the bottom of the tube. The 50 µL of solution was pipette mixed, then 1 µL of the solution was spread on a microscope slide. The slide was allowed to dry for 5 minutes, then pure ethanol (Merck Cat. #1085430250) was added to the surface of the slide to fix the cells. The slide was visually inspected at 40x magnification to ensure that ≥ 95% of the isolated cells were sperm cells. The sperm were then lysed by adding 100 mg of 0.2 mm stainless steel beads (Next Advance Cat. #SSB02) to a BeadBug homogeniser tube (Merck Cat. #Z742479), then transferring the remaining ~49 µL sperm solution to the homogeniser tube. A sperm lysis buffer was prepared by adding the following to an empty 1.5 mL tube: 497.5 µL Qiagen Buffer RLT (Qiagen Cat. #79216) and 2.5 µL of 0.5 M Bond-Breaker TCEP Solution (Thermo Scientific Cat. #77720). 500 µL of sperm lysis buffer was added to the homogeniser tube containing the sperm and beads, then the sample was homogenised in a BeadBug 3 Benchtop Microtube Homogeniser (Pathtech Cat. #032-D1030-E-AU) set to 2800 RPM, using 5 x 60 second cycles with 30 second rests between each cycle to allow for heat dissipation.

Sperm DNA extraction was performed with a modified QIAamp DNA Mini Kit (Qiagen Cat. #51304) protocol. Briefly, 500 µL of buffer AL (QIAamp DNA Mini Kit) was added to the homogenised sample and mixed by vortexing, then 500 µL of pure ethanol (Merck Cat. #1085430250) was added and mixed by vortexing. 600 µL of the solution was applied to a QIAamp Mini spin column in a 2 mL collection tube, and centrifuged at 6000 x g for 1 minute. The flow-through was discarded and this process was repeated until the entire solution had been spun through the column. The remaining spin and wash steps (following application to the column) were performed as per the kit manual, and the elution was performed into a 1.5 mL LoBind tube (Eppendorf Cat. #30108051) with 100 µL of low-Tris (10mM Tris-HCl pH 8) buffer. An RNase master mix was then prepared by adding the following to an empty 1.5 mL tube: 30 µL PBS (10X) pH 7.4 (Thermo Scientific Cat. #70011044), 1 µL RNase A (Qiagen Cat. #19101), and 19 µL nuclease-free water (NEB Cat. #B1500L). 20 µL of RNase master mix was added to the eluted DNA and pipette mixed, then the solution was incubated at room temperature for 5 minutes. The DNA was then purified with a 0.8X SPRIselect (Beckman Coulter Cat. #B23318) bead cleanup and eluted into 35 µL of LTE buffer (10mM Tris-HCl pH 8, 0.1mM EDTA). Eluted DNA was stored at -20°C in a 1.5 mL LoBind tube (Eppendorf Cat. #30108051) until use.

#### Library preparation and sequencing

For the datasets presented in this publication, two versions of the library preparation protocol were used. Version 13 was performed as per Basic Protocol 1 and Basic Protocol 2 from Phie *et al*. (2026).

Version 10 was an earlier iteration of the same protocol. The primary differences from Version 13 were:

a) Nuclease-free water was used instead of LTE buffer for steps 2, 4, 11, 20, 60, 61 and 65 of Basic Protocol 1
b) Samples were stored at ambient temperature instead of -20°C prior to sequencing.

Protocol version 10 was superseded due to oxidative damage signatures present in some samples.

All datasets presented in this publication were sequenced as per Phie *et al*. (2026).

#### Bioinformatics pipeline

All datasets were processed using SomaticCODEC v5.0.0 (Johnstone, Phie and Fraser, 2026). The workflow is summarised in Supplementary Figure 3.

SomaticCODEC uses a matched sample design. An EX (experimental) sample is used to call somatic variants, and an MS (matched) sample is used to identify germline variants and improve variant calling. Alternative terminology used in the literature to describe similar designs includes case/control and tumour/normal.

MS reads are quality trimmed, filtered, aligned, annotated, and deduplicated by UMI family and alignment position. A pileup is performed on MS reads to identify potential germline variant positions. Low-depth positions (<40× by default) are used to generate a low-depth mask. These masks are combined with masks for difficult or repetitive regions and high-frequency germline variants (allele frequency ≥0.1 by default). The combined mask is inverted to define an "include BED" specifying positions eligible for somatic variant calling.

EX reads are demultiplexed, quality trimmed, filtered, and aligned. Reads are then filtered to remove non-properly paired reads and non-primary alignments, annotated, and grouped by UMI family and alignment position. Duplex consensus reads are generated, realigned, annotated, and filtered to remove reads with low mapping quality (MAPQ <50 by default). Somatic SNVs are then called by comparing the duplex consensus reads to GRCh38, at positions defined by the include BED.

Tool versions and dependencies are specified in the environment.yml file in the project repository.

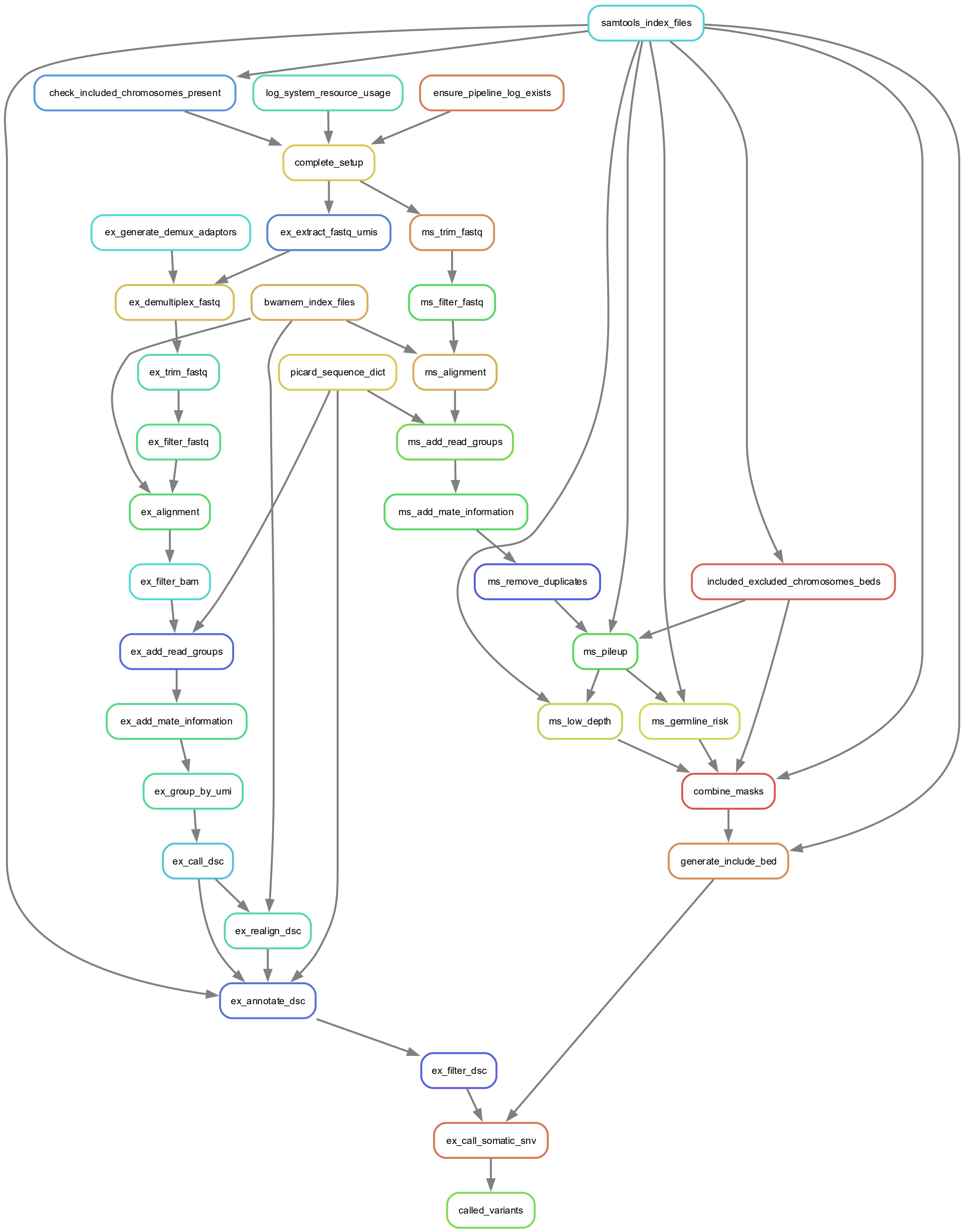


**Supplementary Figure 3 – Directed Acyclic Graph (DAG) for the SomaticCODEC pipeline**
The DAG represents the hierarchy of Snakemake jobs in the SomaticCODEC pipeline. Each node represents a rule, and arrows connect each rule to its dependencies and successors. The two main branches of the pipeline are the matched sample (MS) rules, which create a mask for potential germline variants and difficult or repetitive regions, and the experimental sample (EX) rules, which generate duplex consensus reads and call somatic variants at unmasked positions.

#### Metrics reported in publication

##### Linearity

Sperm and whole blood were obtained from a 43 year old male. N = 16 mixtures were created ranging with even spacing from 0% blood DNA to 100% blood DNA. Libraries were prepared as per protocol version 10. Association between percentage blood and SNV burden was estimated with simple linear regression. One-sided confidence intervals were used as only performance below the specified threshold was considered relevant.

##### Intra-batch precision

A buffy coat remnant sample was obtained from a 70 year old female donor. This sample was split into n = 12 aliquots, with each sample independently processed from DNA extraction to variant calling. Libraries were prepared as per protocol version 13. Coefficient of variation (CV) and normalised interquartile range (norm-IQR) were calculated from SNV burden reported for each sample. One-sided confidence intervals were used as only performance below the specified threshold was considered relevant.

##### Buffy coat SNV burden vs age

Buffy coat remnant samples were collected from 60 independent donors between the ages of 20 and 81. Individuals were ineligible to donate samples if they had high or low blood pressure, high or low temperature, cold or flu symptoms, tattoos in the last 4 months, were pregnant, had noted heart conditions, low iron/haemoglobin levels, certain sexual activities within the past 3 months, injected recreational drugs in the past 5 years, or visited specific countries in the last 4 months.

Power was estimated using Monte Carlo simulation, varying the number of batches and model parameters (slope, intercept, residual error, inter-batch variability, and attrition rate) across predefined distributions informed by pilot data and prior literature. A minimum power of 90% was targeted to detect a conservative estimate of the association between SNV burden and age. Based on these simulations, three batches of 12 donors (36 total) were selected.

As donors were recruited over time, batches were selected from the available pool using a greedy nearest-neighbour algorithm to optimise for a uniform age distribution and balanced sex distribution both within each batch and across the study.

Libraries were prepared according to protocol version 10. The association between SNV burden and donor age was estimated using simple linear regression. The slope was compared to buffy coat data from Bae *et al*. (2023) and granulocyte data from Abascal *et al*. (2021). Slopes for SNV burden versus donor age from previous studies were recalculated from raw data using simple linear regression to ensure direct comparability with SomaticCODEC.

##### Sperm SNV burden

Sperm was obtained from 3 male donors aged 25, 28 and 29 years old respectively. Libraries were prepared as per protocol version 13. SNV burden was compared to external studies using ANOVA and Tukey's HSD post hoc.

**gnomAD overlap**

The results reported in this publication were derived from the dataset described above for Buffy coat SNV burden vs age (n = 36 buffy coat). For each sample, overlap with the gnomAD v4 database, (AF > $1 \times 10^{-3}$) was computed as a percentage of called variants present in both.

**Read position bias**

Buffy coat remnant samples (n=9) were derived from donors spanning the ages 24-69 years old. Libraries were prepared as per protocol version 10. For each called somatic SNV, duplex reads overlapping the SNV were identified, and the position of the SNV in these reads was recorded as a percentile of the read length. The read length percentiles for each SNV were then used to calculate an empirical cumulative distribution function for SNVs over the read length, which was compared to a uniform distribution. The maximum absolute deviation (MAXD) from the uniform distribution was computed for each sample. Across all samples, the median MAXD was reported.

**Genomic distribution index**

The results reported in this publication were derived from the dataset described above for Buffy coat SNV burden vs age (n = 36 buffy coat). The position of each called somatic SNV on its respective chromosome was calculated as a percentage of chromosome length (as determined from the reference genome index file). The mean absolute deviation (MAD) from a uniform distribution was calculated for each chromosome. The reported *genomic distribution index* was defined as the MAD for the $80^{th}$ percentile worst chromosome.

**Duplex disagreement rate**

The results reported in this publication were derived from the dataset described above for Buffy coat SNV burden vs age (n = 36 buffy coat). The duplex disagreement metric was calculated as the proportion of eligible duplex bases where Watson and Crick strands disagreed. Bases were considered eligible for assessment in this metric if they passed all other variant call eligibility filters, except for the requirement that the Watson and Crick strands agree.

**Trinucleotide context**

The results reported in this publication were derived from the dataset described above for intra-batch precision (n = 12 buffy coat from same donor). This dataset was selected for this analysis as it used library preparation protocol version 13, that had reduced oxidative damage which impacted the trinucleotide signature. The limitation of this dataset is that it is derived from a single donor.

The trinucleotide context of each sample was compared to a consensus context for granulocytes derived from Abascal *et al*. (2021), by computing cosine similarity.

##### Other metrics

For metrics reported in Supplementary Table 1 that are not described in the main text, refer to the GitHub repository for details on their implementation. The authors are available to provide further information on the rationale and limitations of these metrics upon reasonable request.

#### Data analysis

All statistical analyses were performed in R v4.3 (R Core Team, 2021) using the tidyverse v2.0.0 (Wickham *et al.*, 2019).

Unless otherwise stated, confidence intervals were calculated as follows. Confidence intervals for medians were estimated using a nonparametric order statistic approach based on the binomial distribution. All other confidence intervals were estimated using bootstrap resampling.

All figures were generated in R using ggplot2 v4.0.0 (Wickham, 2016) and cowplot v1.2.0 (Wilke, 2025).

#### References

Abascal, F. *et al*. (2021) "Somatic mutation landscapes at single-molecule resolution," *Nature*, 593(7859), pp. 405–410. Available at: https://doi.org/10.1038/s41586-021-03477-4.

Bae, J.H. *et al*. (2023) "Single duplex DNA sequencing with CODEC detects mutations with high sensitivity," *Nature Genetics*, 55(5), pp. 871–879. Available at: https://doi.org/10.1038/s41588-023-01376-0.

Johnstone, J.N., Phie, J. and Fraser, C. (2026) *SomaticCODEC*. Systematic Medicine. Available at: https://github.com/systematicmedicine/SomaticCODEC.

Phie, J. *et al*. (2026) "CODEC Library Preparation From Genomic DNA," *Under review* [Preprint].

R Core Team (2021) *R: A language and environment for statistical computing*. Vienna, Austria: R Foundation for Statistical Computing. Available at: https://www.R-project.org/.

Shoag, J.E. *et al*. (2025) "Direct measurement of the male germline mutation rate in individuals using sequential sperm samples," *Nature Communications*, 16(1), p. 2546. Available at: https://doi.org/10.1038/s41467-025-57507-0.

Wickham, H. (2016) *ggplot2: Elegant Graphics for Data Analysis*. New York: Springer-Verlag. Available at: https://ggplot2.tidyverse.org/.

Wickham, H. *et al.* (2019) “Welcome to the Tidyverse,” *Journal of Open Source Software*, 4(43), p. 1686. Available at: https://doi.org/10.21105/joss.01686.

Wilke, C.O. (2025) *cowplot: Streamlined Plot Theme and Plot Annotations for “ggplot2.”* Available at: https://wilkelab.org/cowplot/.
